# Systematic elucidation of independently modulated genes in *Lactobacillus plantarum* reveals a trade-off between secondary and primary metabolism

**DOI:** 10.1101/2023.11.03.565434

**Authors:** Sizhe Qiu, Yidi Huang, Shishun Liang, Hong Zeng, Aidong Yang

## Abstract

*Lactobacillus plantarum* is a probiotic bacteria widely used in food and health industries, but its gene regulatory information is limited in existing databases, which impedes the research of its physiology and its applications. To obtain a better understanding of the transcriptional regulatory network of *L. plantarum*, independent component analysis (ICA) of its transcriptomes was used to derive 45 sets of independently modulated genes (iModulons). Those iModulons were annotated for associated transcription factors (TFs) and functional pathways, and active iModulons in response to different growth conditions were identified and characterized in detail. Eventually, the analysis of iModulon activities reveals a trade-off between regulatory activities of secondary and primary metabolism in *L. plantarum*.

## 1. Introduction

The transcriptional regulatory network (TRN) of a bacterium consists of all regulatory interactions between its transcription factors (TFs) and genes [1]. TFs, also referred to as sequence-specific DNA-binding factors, sense external signals and then bind to promoter regions of operons to regulate gene expression levels [2]. To identify regulatory interactions between TFs and genes, the most commonly used experimental method is chromatin immunoprecipitation followed by sequencing (CHiP-seq) [3]. In CHiP-seq, antibodies are used to select TF proteins, and then DNA bound to TF proteins will be purified. DNA sequencing for the DNA-TF protein complex will determine the binding site on the genome. A group of genes with binding sites of the same TF are considered as a regulon. However, the drawbacks of CHiP-seq lie in its high cost, time-intensive nature, and challenges in capturing the diverse growth conditions of bacteria [4].

In recent years, many computational methods of *in-silico* reconstruction of TRN have been developed, such as coexpression network analysis [5] or supervised learning-based methods (e.g., GENIE3 [6]). One of the most popular methods to reconstruct TRN is using independent component analysis (ICA) to decompose the gene expression matrix, which consists of transcriptomic data of different samples, into sets of independently modulated genes, called iModulons (IMs) [7]. Apart from derived IMs, ICA can also quantify IM activities in different samples. Unlike CHiP-seq being a ‘bottom-up’ method, ICA follows a ‘top-down’ approach. ICA has been extensively applied to study and improve the understanding of many bacteria’s TRNs. For example, ICA of *Vibrio natriegens* transcriptomes unveils the genetic basis of its natural competency [8]. ICA has also been used to discover therapeutic strategies for *Streptococcus pyogenes* by identifying carbon sources that control the expression of hemolytic toxins [9].

*Lactobacillus plantarum* is a gram-positive lactic acid bacterium that can be found in diverse ecological niches [10]. It has been widely used in food and health industries. For instance, it is the major bacterium involved in the fermentation of mozzarella cheese [11]; *L. plantarum*-derived exopolysaccharides (EPSs) have various probiotic effects [12] and anticancer properties [13]. Due to the importance of *L. plantarum* in different biological processes, such as dairy product fermentation, its gene expression regulation has received interest in several studies. For example, Jung and Lee, 2020 identified differentially expressed genes when *L. plantarum* was in the acidic condition [14]. Unlike most studies focusing on single regulatory genes, Wels *et al.*, 2011 reconstructed the gene regulatory network of *L. plantarum* on the basis of correlations between gene expression levels and conserved regulatory motifs [15]. Nonetheless, the regulon information of *L. plantarum* in RegPrecise [16] only recorded 47 regulons and 210 TF binding sites, in contrast to 624 and 943 TF binding sites recorded for *Bacillus subtilis* and *Escherichia coli*, respectively. The lack of gene regulatory information hinders the study of *L. plantarum*’s physiology and rational engineering of this bacteria.

Considering the value of *L. plantarum* in industry and research as well as the limited understanding of its TRN, this study managed to infer undiscovered regulatory interactions using ICA decomposition of the gene expression matrix, and to further investigate how *L. plantarum* respond to different growth conditions (e.g., acid stress). Moreover, this study, through the analysis of IM activities, explored the growth strategy of *L. plantarum*, in terms of how it balances different biological processes (e.g., energy generation, carbohydrate metabolism, stress responses).

## 2. Methods

### 2.1 Data acquisition and preprocessing

The transcriptomic data used in the study were obtained from 4 independent studies that included various experimental conditions: response to pH decrease from 6.2 to 5.0 [14], treatment with N-3-oxododecanoyl homoserine lactone (a quorum sensing molecule) [17], contrasting habitats (e.g., bee extract) [18], and change of carbon sources [19]. The metadata of sample conditions can be found in **SI, Table S1**. In the data from the selected 4 studies, genes were all annotated based on the genome assembly of *L. plantarum* WCFS1 (ASM20385v3) [20]. All transcriptomic sequencing reads were normalized as RPKM (Reads Per Kilobase Million). Then, all samples were merged as a compendium of transcriptomic data (100 samples*3000 genes). Before independent component analysis (ICA) was undertaken, the merged dataset was first log-transformed, and then centered by subtracting the expression levels of the reference condition (i.e., wt_pH6.2 in **SI, Table S1**). The data quality was demonstrated by the higher Pearson correlation coefficients (PCCs) between replicates than PCCs between non-replicates [21] (**Figure 1A**).

**Figure 1.**
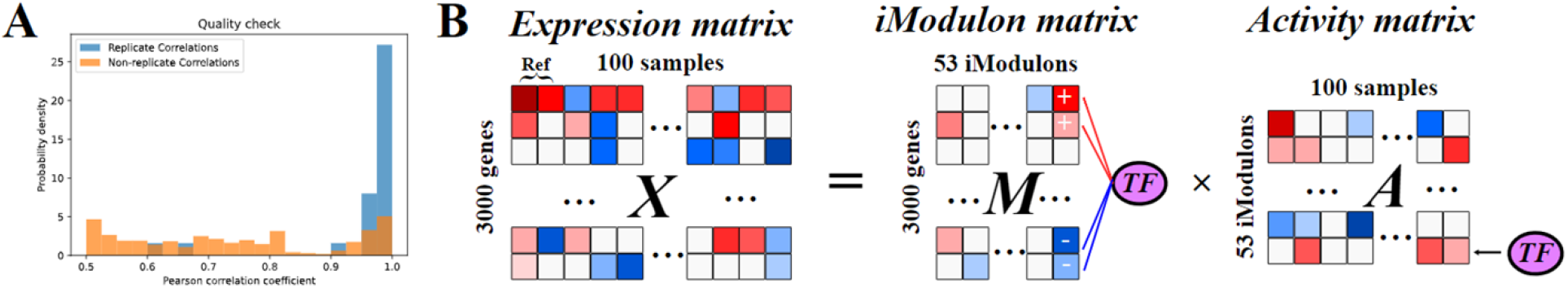
ICA decomposes the compendium of transcriptomic data to 45 nonempty iModulons. (A) Quality check of transcriptomic data with PCCs. Blue: replicate correlations; Yellow: non-replicate correlations. (B) Schematic illustration of ICA applied to the gene expression matrix.

### 2.2 Determination of iModulons

ICA decomposition of the merged dataset (i.e., the expression matrix, 3000 genes * 100 samples) was conducted using scripts in precise-db (https://github.com/SBRG/precise-db) [22]. The FastICA algorithm in Scikit-Learn (v0.20.3) [23] was used to calculate independent components with 100 iterations with a tolerance of 10^―7^, *log*(*cosh(x)*) as the contrast function, and parallel search algorithm. The OptICA method was used to determine the optimal number of independent components [24]. The output of ICA were the iModulon matrix (M matrix, 3000 genes * 53 IMs) and Activity matrix (A matrix, 53 IMs * 100 samples) (**Figure 1B**). The M and A matrices can be found in https://github.com/SizheQiu/LPiModulons/tree/main/data/IMdata.

Gene weights in each column (for the corresponding IM) of the M matrix were used to determine each gene’s IM membership. The threshold of gene weight absolute values for each IM was computed based on D’Agostino’s *K*^2^test using the PyModulon package (https://github.com/SBRG/pymodulon) [7]. The default *K*^2^-statistic cutoff of 550 was used. The genes with weight absolute values above the threshold were the member genes of the IM. Before annotation, IMs were labeled as IM-1 to 53.

### 2.3 Annotation of iModulons via regulon enrichment analysis

Regulons of *Lactobacillus plantarum WCFS1* were obtained from RegPrecise [16]. IMs that overlap with regulons were annotated via regulon enrichment analysis. The set of genes in each IM was compared to each regulon using the two-sided Fisher’s exact test (False Discovery Rate (FDR) < 10^―5^) [7]. After regulon enrichments were computed for IMs, regulatory annotations were manually determined based on the venn diagrams of IMs and regulons (s**ee SI, Figure S1**). In addition to IMs associated with only one regulon (e.g., PyrR IM (IM-36)), there were two different annotation expressions for combined regulon enrichments: intersection (+) and union (/). If a specific combinatorial regulation (genes controlled by multiple regulators) was observed in the venn diagram of the IM and enriched regulons, then the IM was annotated with regulators linked by “+” (e.g., MalR+MdxR IM (IM-47)). Otherwise, “/” was used (e.g., ArgR/MleR IM (IM-26)).

### 2.4 Annotation of iModulons via motif comparison

IMs that do not overlap with known regulons were annotated via motif discovery and motif comparison. If a coding gene’s 200 bp upstream region does not overlap with another gene [25] and BDGP Neural Network Promoter Prediction [26] predicted this region to be a possible promoter (probability score > 0.8), then this 200 bp upstream region was used to search for sequence motifs using MEME [27]. Motif comparison by TOMTOM [28] then determined the most possible TF based on the similarity of found motifs and TF binding site motifs in databases (e.g., RegTransBase [29]). The p-value and E-value thresholds set in TOMTOM were 0.05 and 10. To further validate whether genes in the IM are regulated by the found TF, PCCs of the expression levels of the TF gene and IM genes were computed. If the gene had significant correlations (p-value < 0.05) with most genes in the IM, then the TF would be used to annotate the IM.

## 3. Results

### 3.1 Regulatory and functional annotations of identified iModulons

The derived 53 IMs account for 85% explained variance of the gene expression matrix. In each IM, genes with absolute values of weights higher than the threshold are determined as IM member genes (see Methods, section 2.2). The details of IM member genes can be found in https://github.com/SizheQiu/LPiModulons/tree/main/data/IMdata/ as IM_genes.csv. Among 53 IMs, 45 are nonempty and most IMs’ sizes are within 20 (Figure 2A). Only 17% IM member genes overlap with genes in known regulons (Figure 2B), and hence only 13 IMs could be annotated via regulon enrichment (Figure 2C). The details of regulatory annotations can be found in https://github.com/SizheQiu/LPiModulons/blob/main/data/IMdata/IM_annotation.csv.

**Figure 2.**
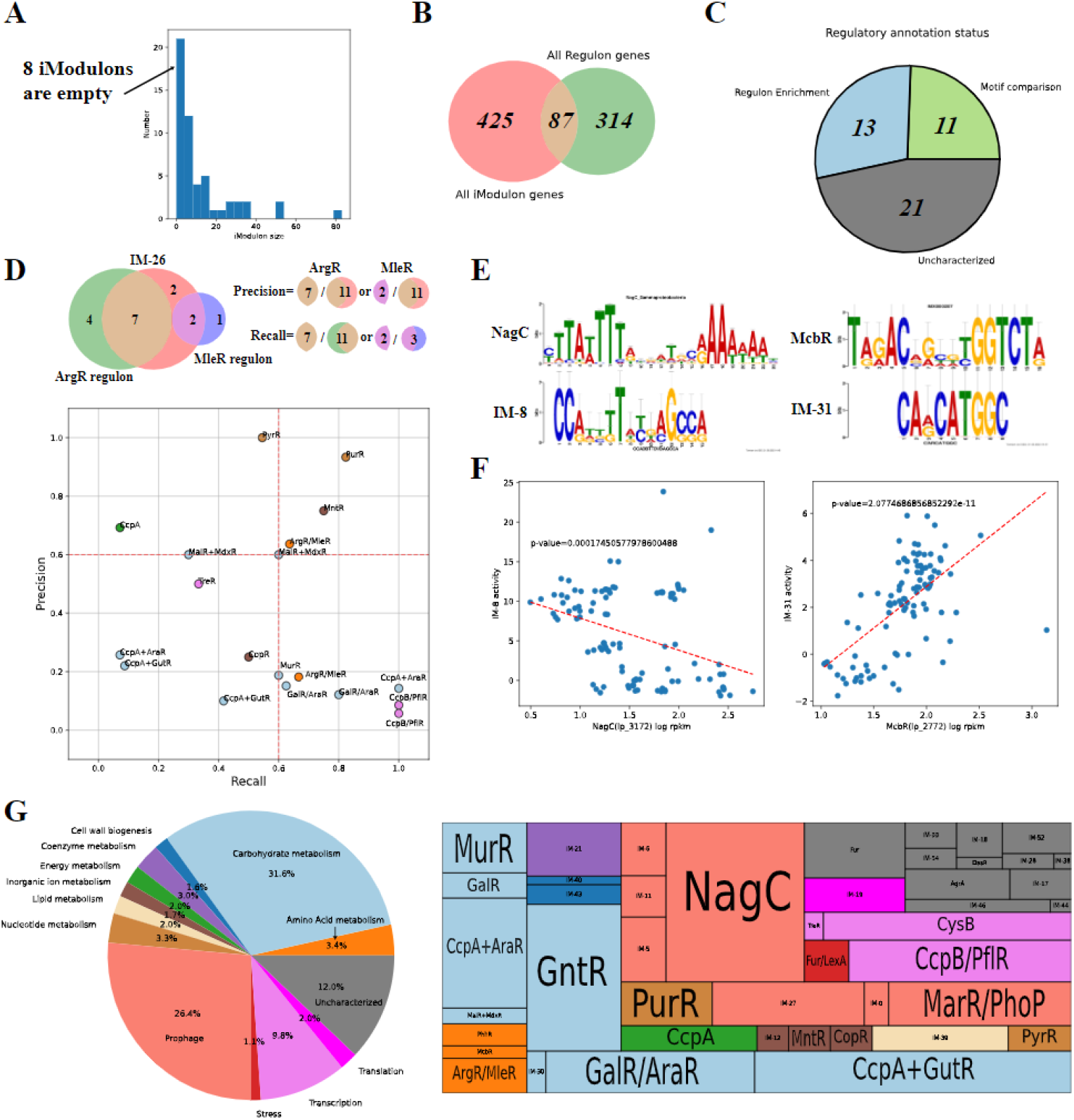
Regulatory and functional pathway annotations of IMs. (A) The histogram of IM sizes, 45 out of 53 IMs are nonempty. (B) The venn diagram of all IM genes and regulon genes. 87 genes in IMs are contained in known regulons. (C) The pie chart of regulatory annotation status. Blue: regulon enrichment; Green: motif comparison; Grey: uncharacterized. (D) Recall and precision of IMs with matched regulons. (E) Motif comparison of IM-8 and IM-31. (F) The significant correlations between IM activities and gene expression levels of associated TFs identified via motif comparison for IM-8 and IM-31 (p-value < 0.05). (G) The pie chart and treemap of functional annotations of IMs, the size of each fraction is scaled with the IM size.

For the 13 IMs annotated with enriched regulons, most of them have either high recall or high precision (cutoff = 0.6) (Figure 2D). Venn diagrams showing regulon enrichments in IMs are provided in **SI, Figure S1**. High recall means that the overlap (of IM and regulon) has high coverage of the regulon, while high precision means that the overlap has high coverage of the IM. IMs with low recall and low precision are considered to be incompletely matched with regulons, but that does not necessarily mean the IM’s regulatory annotation is inaccurate. For example, the remaining 3 genes in CopR IM that are not included by the current CopR regulon of *Lactobacillus plantarum WCFS1* are lp_3055(copA), lp_3057(copper-binding protein) and lp_3058(copper-binding protein), but they are included by the CopR regulon of other closely related lactic acid bacteria (e.g., *Lactococcus lactis subsp. lactis Il1403*) [30].Therefore, the low recall and precision are sometimes resulted by the incompleteness of currently known regulons.

In addition to IMs associated with regulons, there are 11 IMs annotated via motif search and comparison (Figure 2C, Figure S2). Two representative examples are NagC IM (IM-8) and McbR IM (IM-31) (Figure 2E). Their regulatory annotations are validated by significant correlations between expression levels of TF genes and IM activities (Figure 2F). The remaining 21 IMs (Figure 2C) cannot be annotated via motif search and comparison either because the IM does not contain multiple possible promoter sequences for motif search (e.g., IM-19) or TOMTOM (Methods, section 2.4) fails to find a TF binding site motif with a high similarity to the found motif (e.g., IM-6).

IMs were also annotated with enriched functional pathways (**see SI, 3.1**), and the details of functional annotations can be found in https://github.com/SizheQiu/LPiModulons/blob/main/data/IMdata/IM_annotation.csv. Apart from the uncharacterised group, 3 dominant functions of derived IMs are carbohydrate metabolism, prophage proteins and transcription (Figure 2G). Fur/LexA IM (IM-1) was functionally annotated as “Stress”, as LexA has already been found as a TF for stress response [31]. IM-19 was annotated as ‘Translation’, because genes in IM-19 were all ribosomal genes (e.g., rplV (lp_1039), large ribosomal subunit protein uL22). 12% (scaled with IM sizes) of IMs are uncharacterized in functional annotation due to the lack of enriched functional pathways.

### 3.2 Comparison between iModulons and regulons

The difference between IMs and regulons can provide undiscovered regulatory information. Regulon enrichments of some IMs show combinatorial regulations of multiple TFs, such as MalR+MdxR IM. Based on the genomic organization, 6 genes in the region between 151222bp and 158185bp belong to the same operon (Figure 3A). While mdxE (lp_0175), mdxG (lp_0177) and lp_0178 are already included by both MalR and MdxR regulons, MalR+MdxR IM also captures the combinatorial regulatory signals for malS (lp_0179) and msmX (lp_0180), which share the same promoter with genes in the overlap of MalR and MdxR regulons. All genes in MalR+MdxR IM are involved in maltose/maltodextrin metabolism, which is the biological process regulated by MalR and MdxR [32][31].

**Figure 3.**
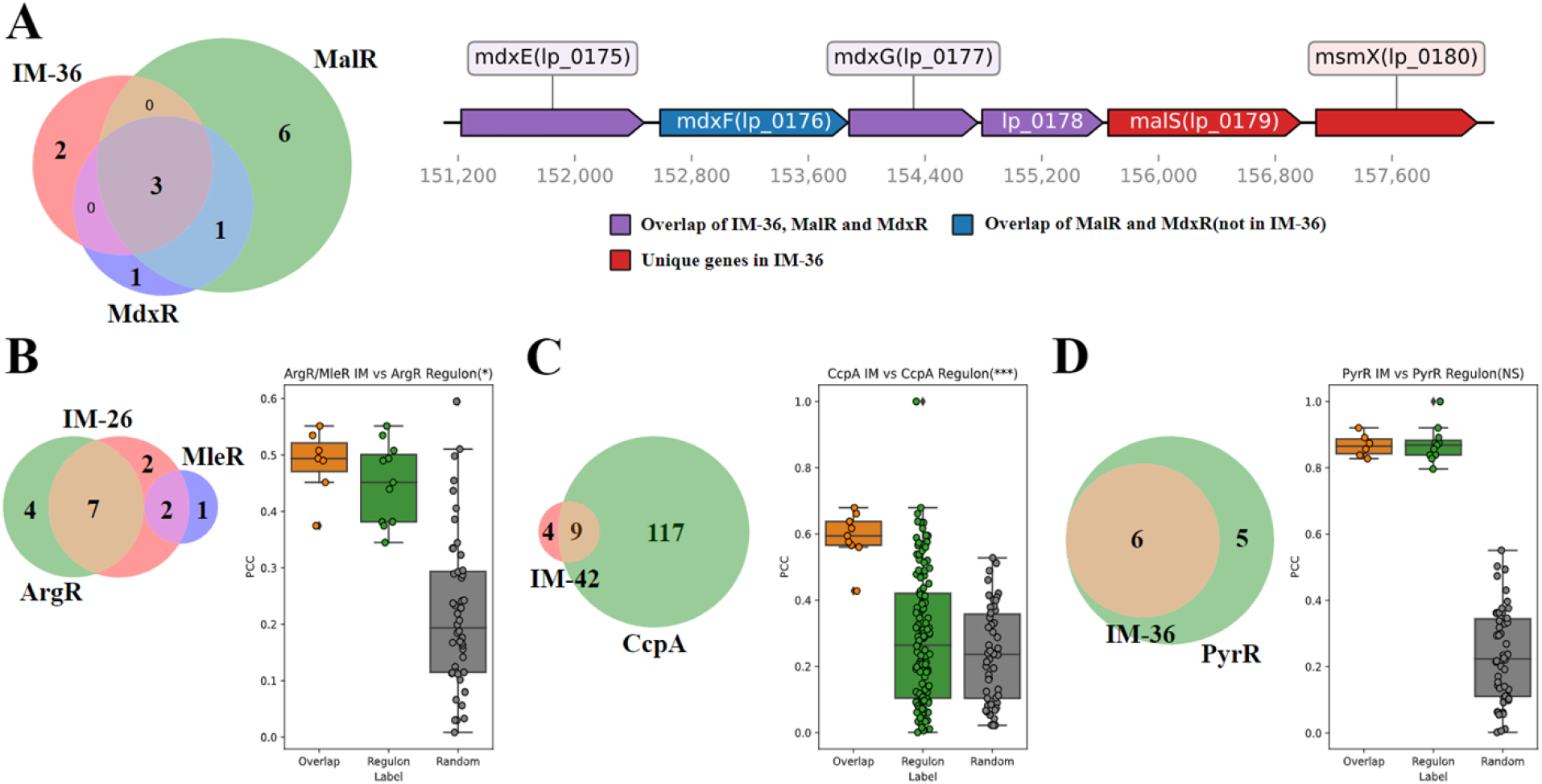
Comparison between IMs and regulons of *Lactobacillus plantarum*. (A) Left: venn diagram; Right: Genomic organization of genes in MalR+MdxR IM (IM-47). (B-D) Comparison of PCCs of the gene expression levels of TF gene and genes in the overlap of IM and regulon (orange), regulon (green) and randomly sampled genes (gray) for (B) ArgR/MelR, (C) CcpA, and (D) PyrR IMs.

IMs also have the ability to identify genes with strong regulatory interactions with TFs from known regulons. For example, the Pearson correlation coefficients (PCCs) between TF genes and genes in the overlap (of the IM and regulon) exhibit higher distributions compared to those of genes in the regulon for ArgR and CcpA (**Figure 3BC**). Nevertheless, the overlap does not always show stronger regulatory interactions. For example, genes in PyrR IM do not have significantly higher PCCs with the PyrR gene than with the genes in PyrR regulon (Figure 3D).

### 3.3 Active iModulons in response to different growth conditions

In addition to the M matrix, the A matrix is another output of ICA decomposition, which reveals IM activities of *L. plantarum* under different growth conditions. In response to acid stress (in terms of pH decrease), 4 active IMs are observed: Fur/LexA IM, CopR IM, McbR IM and PyrR IM (Figure 4A). The gene expression levels of Fur (lp_3247) and LexA (lp_2063) both decrease with the decrease of pH, though the trends over three pH values are not consistently decreasing (Figure 4B). Genes in Fur/LexA IM are related to the biosynthesis of exopolysaccharide (EPS), an important secondary metabolite [33], including lp_0302 (extracellular transglycosylase), lp_0304 (extracellular transglycosylase), lp_2809 (extracellular protein of unknown function), lp_2810 (glycosyl hydrolase, family 25), lp_2845 (extracellular transglycosylase, with LysM peptidoglycan binding domain), lp_3014 (extracellular transglycosylase, with LysM peptidoglycan binding domain) and lp_3050 (extracellular transglycosylase, membrane-bound). Oppositely, the gene expression levels of CopR (lp_3365), McbR (lp_2772) and PyrR (lp_2704) increase with the decrease of pH (Figure 4CDE). CopR, McbR and PyrR regulate copper homeostasis, amino acid metabolism and pyrimidine metabolism, respectively.

**Figure 4.**
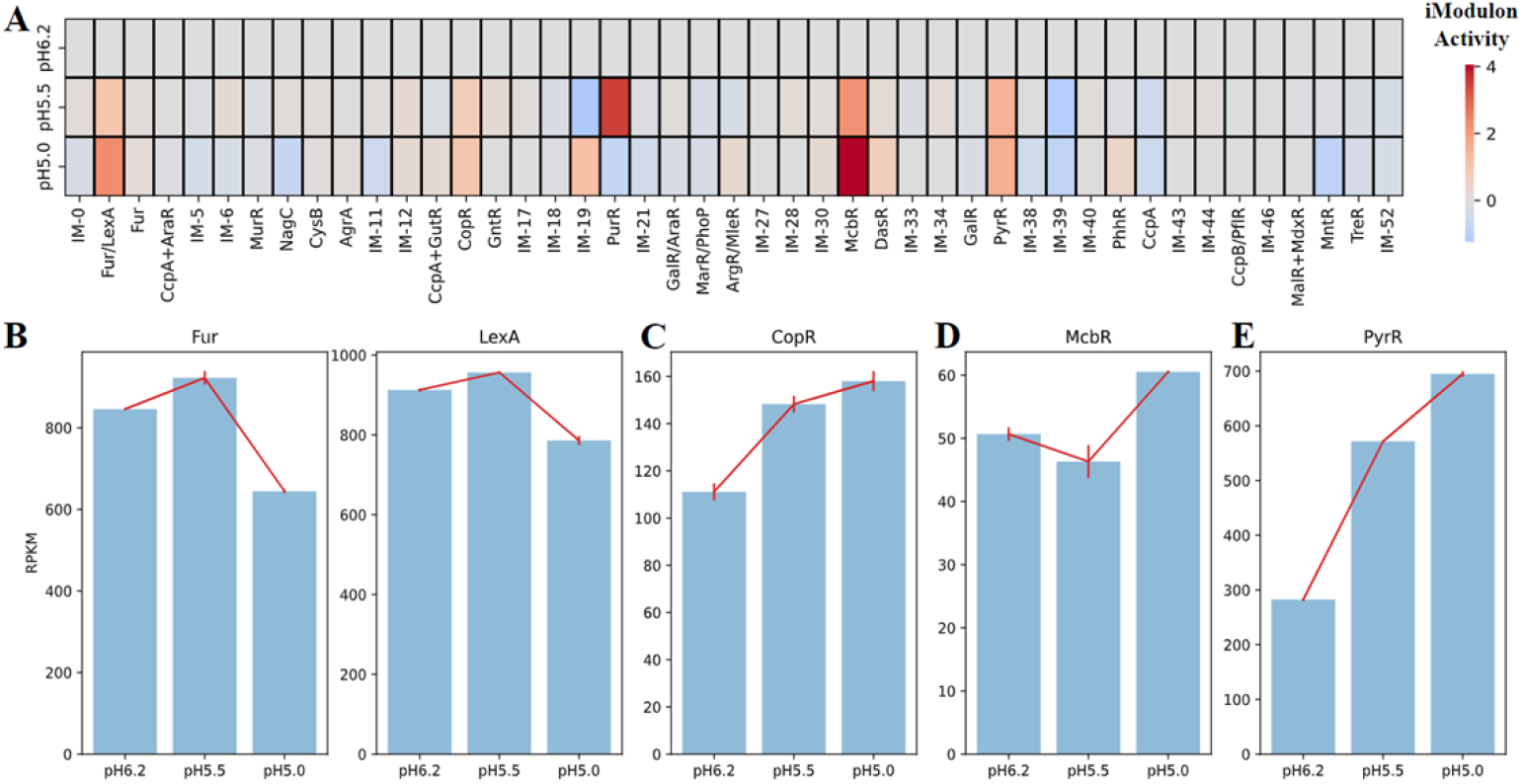
Identification of active IMs under the acidic condition. (A) The heatmap of IM activities at pH=6.2, 5.5 and 5.0. (B) The expression levels of Fur (lp_3247) and LexA (lp_2063) at different pH values. (C) The expression levels of CopR (lp_3365) at different pH values. (D) The expression levels of McbR (lp_2772) at different pH values. (E) The expression levels of PyrR (lp_2704) at different pH values.

To further characterize acid-active IMs, regulatory networks are reconstructed as weighted correlation networks, and genomic organizations of genes in those IMs are further investigated. Fur/LexA IM, based on gene locations and the weighted correlation network, appear to contain two operons regulated by Fur and LexA separately: lp_0302 and lp_0304 regulated by Fur; lp_2809 and lp_2810 regulated by LexA (**Figure 5AB**). The correlations between Fur and lp_0302, lp_0304 and lp_3014 are all negative, consistent with the previous finding that Fur is a repressor [34] (**Figure 5A**). The correlations between LexA and its regulated genes (i.e., lp_2809, lp_2810 and lp_3050) are positive, indicating that LexA functions as an activator to those genes (**Figure 5A**). For CopR, McbR and PyrR IMs, the correlations between TFs and regulated genes are all positive, suggesting that associated TFs all function as activators (**Figure 5CEG**). Unlike Fur/LexA IM, member genes of those three IMs are mainly in single operons (**Figure 5DFH**).

**Figure 5.**
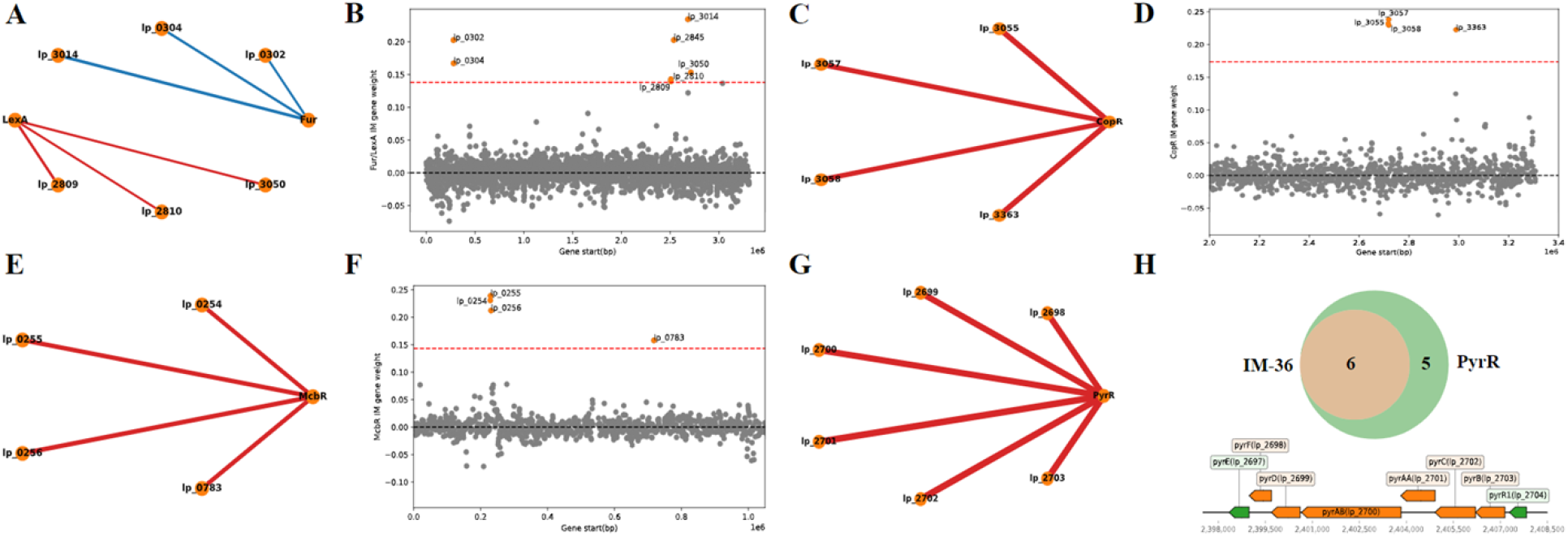
Characterization of genes in acidity-active IMs. (A) The weighted correlation network of Fur, LexA and genes in Fur/LexA IM (IM-1). (B) Gene weights and gene locations of Fur/LexA IM. (C) The weighted correlation network of CopR and genes in CopR IM (IM-15). (D) Gene weights and gene locations of CopR IM. (E) The weighted correlation network of McbR and genes in McbR IM (IM-31). (F) Gene weights and gene locations of McbR IM. (G) The weighted correlation network of PyrR and genes in PyrR IM (IM-36). (H) Genomic organization of genes in PyrR IM (IM-36). Orange: overlap of IM and regulon; Green: genes only in the regulon. Edge weights in weighted correlation networks are scaled to PCCs. Red: positive correlation; Blue: negative correlation; Orange node: gene.

On the other hand, the change of carbon sources can result in transcriptional regulations of carbohydrate metabolism [35], where GntR IM (IM-16) was found to be the most active IM in this study (Figure 6A). Genes in GntR IM mainly encode for the utilization of different carbon sources (e.g., pts9C (lp_0576), uptake of mannose; panD (lp_0579), aspartate 1-decarboxylase) and the biosynthesis of capsular polysaccharide (CPS) in the cell wall (e.g., cps1F (lp_1182), CPS biosynthesis protein CpsC). The biosynthesis of CPS is a part of primary metabolism (cellular biomass formation), different from that of EPS, belonging to secondary metabolism [36]. GntR IM is annotated via motif comparison (Figure S2) due to the lack of regulon information, and hence it is hard to determine which TF in the GntR family regulate genes in this IM. Top 4 GntR TF genes with highest PCCs with activities of GntR IM are lp_2615, lp_2651, lp_3633 and lp_0563 (Figure 6B). The PCCs between TF genes and genes in GntR IM show that lp_2615 and lp_0563 have significant negative correlations with genes in GntR IM, while lp_2651 and lp_3633 have significant positive correlations with genes in GntR IM (Figure 6C), which are consistent with the PCCs (Figure 6B). Possibly, genes in GntR IM are regulated by multiple GntR family TFs.

**Figure 6.**
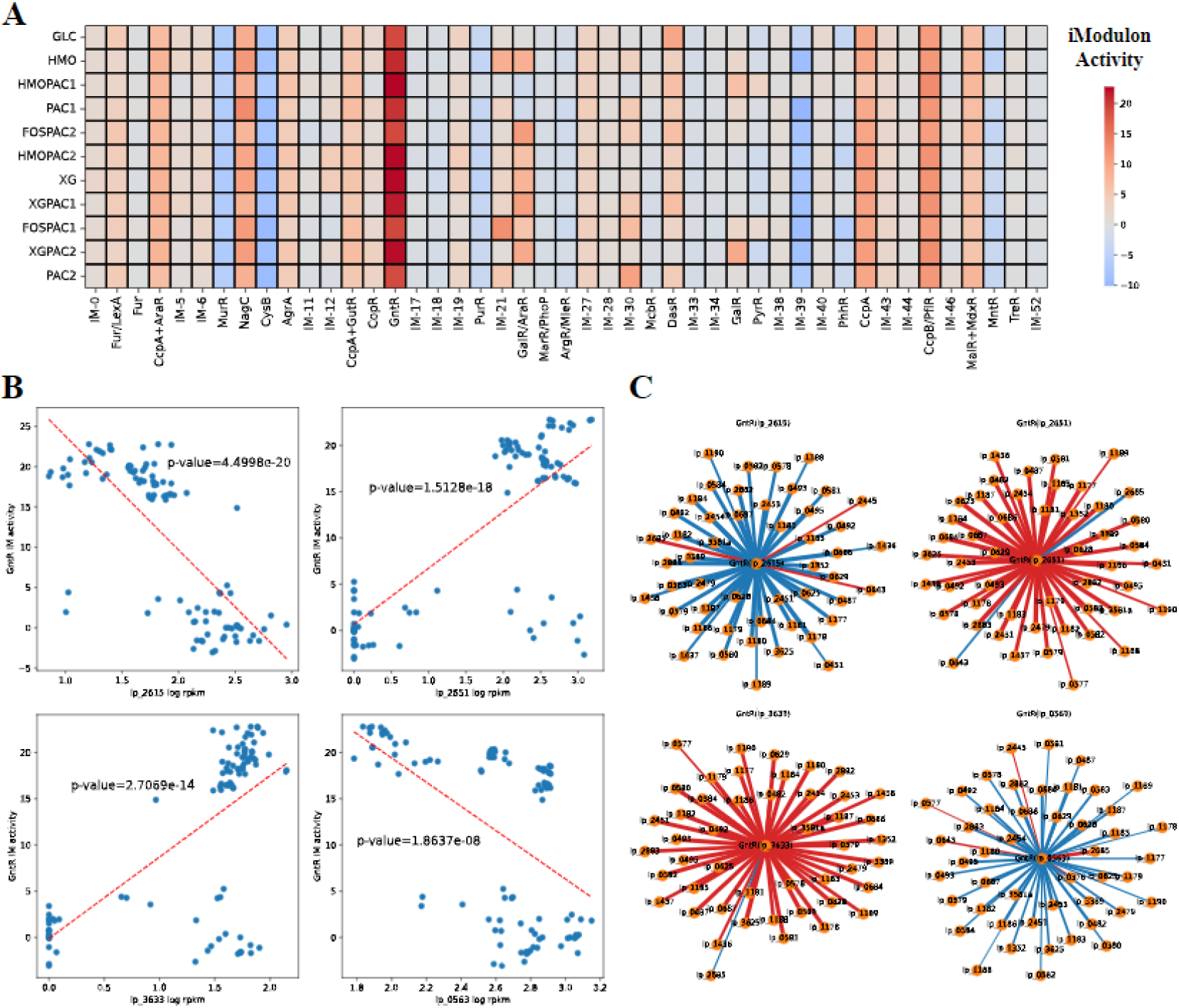
Identification of the most active IM in response to different carbon sources: GntR IM (IM-16). (A) The heatmap of IM activities with different carbon sources. GLC: glucose; HMO: human milk oligosaccharides; PAC1: proanthocyanidin fraction 1; PAC2: proanthocyanidin fraction 2; FOS: fructooligosaccharides; XG: xyloglucans. Detailed information can be found in Özcan *et al.*, 2021 [19]. (B) The correlations between expression levels of 4 GntR family TF genes and GntR IM activities (p-value < 0.05). Red dashed line: linear fit. (C) The weighted correlation networks of 4 GntR family TF genes and genes in GntR IM (p-value < 0.05). Edge weights are scaled to PCCs. Red: positive correlation; Blue: negative correlation; Orange node: gene.

### 3.4 The trade-off between primary and secondary metabolism revealed by iModulon activities

Member genes of IMs derived in this study encode connected reactions in one or several metabolic pathways, and those reactions were visualized as networks (**see SI, 3.2**) to investigate the links between IMs and cellular metabolism (Figure 7). For acid-active IMs identified in section 3.3, genes in McbR IM and PyrR IM encode for the biosynthesis of L-cysteine and uridine monophosphate, respectively (Figure 7AB). EPS biosynthetic reactions encoded by genes in Fur/LexA IM and copper homeostasis encoded by genes in CopR IM are currently not included by model iBT721.

**Figure 7.**
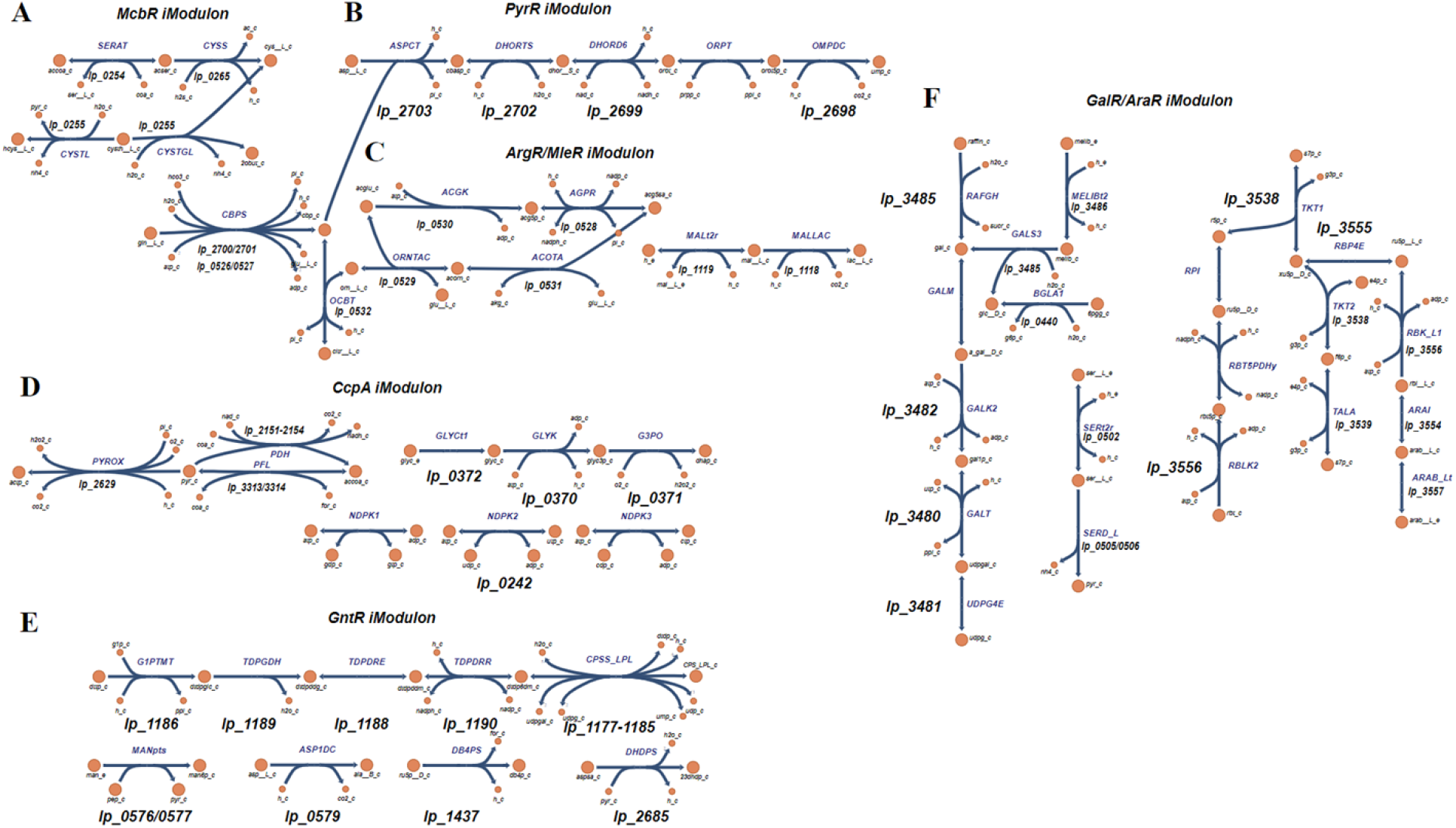
Metabolic pathways encoded by IM member genes. Reaction information (names, associated genes and IMs) can be found in **SI, Table S2**. Reaction abbreviations are adopted from the BIGG database (http://bigg.ucsd.edu/) [37]. (A) McbR IM. (B) PyrR IM. (C) ArgR/MleR IM. (D) CcpA IM. (E) GntR IM. (F) GalR/AraR IM.

Next, 4 representative IMs, namely ArgR/MleR IM, CcpA IM, GntR IM, and GalR/AraR IM, functionally annotated for amino acid metabolism, energy metabolism and carbohydrate metabolism are selected to reconstruct metabolic pathways encoded by their member genes (Figure 2G). ArgR/MleR IM member genes encode for the biosynthesis of N-Acetyl-L-glutamate 5-semialdehyde from L-glutamine (Figure 7C). CcpA IM, as an IM for energy metabolism, contains a part of glycolysis, the conversion of glycerol to dihydroxyacetone phosphate and phosphorylation of nucleosides (Figure 7D). GntR IM member genes mainly encode for CPS biosynthesis, from the activation of monosaccharides to the polymerization as explained in section 3.3 (Figure 7E). Two important carbohydrate metabolic pathways, namely galactose metabolism and pentose phosphate pathway are contained by GalR/AraR IM (Figure 7F).

In contrast to Fur/LexA IM controlling secondary metabolism (EPS biosynthesis induced by acid stress) as shown in section 3.3, ArgR/MleR, CcpA, GntR and GalR/AraR IMs (metabolic pathways visualized in **Figure 7C-F**) regulate primary metabolism. To investigate the relationship between regulatory activities of two branches of cellular metabolism, PCCs were computed for the activities of Fur/LexA IM and 4 IMs for primary metabolism (**Figure 8A-D**). Significant inverse correlations between the activity of Fur/LexA IM and activities of ArgR/MleR IM, CcpA IM, GntR IM and GalR/AraR IM can be observed, suggesting a trade-off between the regulatory activities of secondary and primary metabolisms. *Lactobacillus plantarum* in acidic media (e.g., bee extract (pH=4.7), tomato juice (pH=3.5), **see SI, Table S1**) have higher Fur/LexA IM activities and lower IM activities of the 4 IMs for primary metabolism than those in relatively neutral media (e.g., fecal extract (pH=5.9), **see SI, Table S1**). Therefore, the balance between regulations of EPS biosynthesis and primary metabolism in *Lactobacillus plantarum* appears to significantly depend on the acidity of extracellular environments.

**Figure 8.**
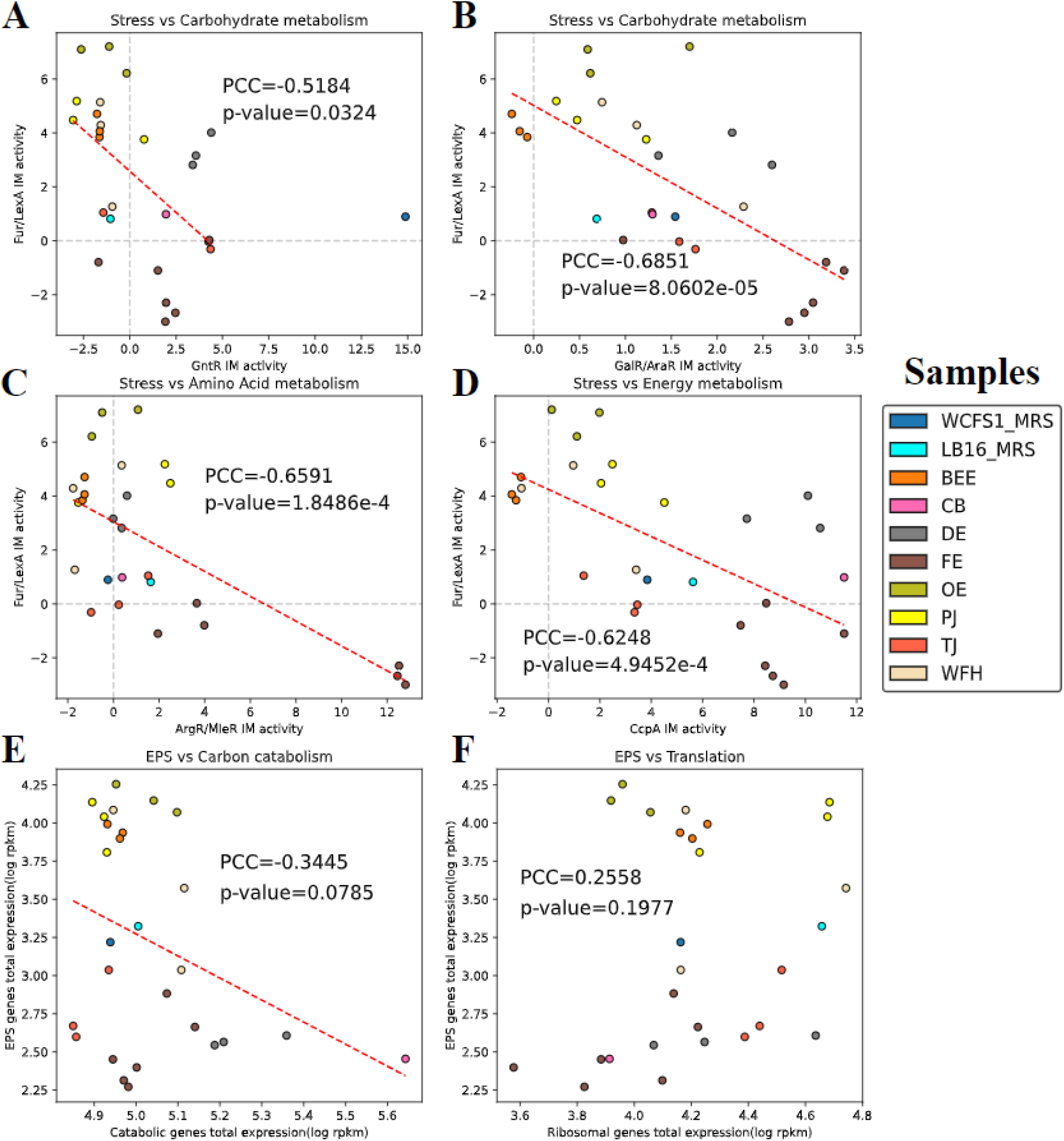
The relationships between secondary and primary metabolisms for *L. plantarum* cultivated in different growth conditions. (A) Fur/LexA IM activity versus GntR IM activity. (B) Fur/LexA IM activity versus GalR/AraR IM activity. (C) Fur/LexA IM activity versus ArgR/MleR IM activity. (D) Fur/LexA IM activity versus CcpA IM activity. (E) The total expression levels (log RPKM) of central catabolic genes and EPS biosynthetic genes (genes in Fur/LexA IM). (F) The total expression levels (log RPKM) of ribosomal genes (genes in Translation IM (IM-19)) and EPS biosynthetic genes. WCFS1_MRS: *L. plantarum* WCFS1 in MRS broth; LB16_MRS: *L. plantarum* LB16 in MRS broth; BEE: Bee extract; CB: Cheese broth; DE: Drosophila sp. extract; FE: Fecal extract; OE: Olive extract; PJ: Pineapple juice; TJ: Tomato juice; WFH: Wheat flour hydrolyzate.

To assess whether a trade-off relationship also exists between gene expression levels (in addition to regulatory activities) of secondary and primary metabolism, PCCs were computed between the total expression levels of genes in Fur/LexA IM (EPS biosynthetic genes) and (i) all glycolytic genes (central carbon catabolism) (**Figure 8E**), and (ii) genes in Translational IM (IM-19, ribosomal genes) (**Figure 8F**). An inverse correlation between gene expression levels of EPS biosynthetic genes and glycolytic genes is also observed (**Figure 8E**), though the correlation is not statistically significant. For EPS biosynthetic genes versus ribosomal genes, there is no inverse correlation between them (**Figure 8F**).

## 4. Discussion

ICA decomposition of *Lactobacillus plantarum* transcriptomes allowed us to identify 45 nonempty IMs, 53.3% of which were annotated with associated TFs via either regulon enrichment analysis (13 IMs) or motif comparison (11 IMs). Annotated IMs revealed several regulatory interactions that have not been reported by known regulons of *L. plantarum*, e.g., malS (lp_0179) and msmX (lp_0180) captured by MalR+MdxR IM (section 3.2), which contributed to the reconstruction of a more complete TRN. Furthermore, the Activity matrix (A matrix) output by ICA decomposition showed the change of regulatory activities of TFs in response to different growth conditions (e.g., acid stress, carbon source switch), leading to the identification and characterization of relevant active IMs (section 3.3). Lastly, the analysis of relationships between IM activities unveiled a trade-off between secondary metabolism (acid stress induced EPS biosynthesis) and primary metabolism in *L. plantarum* (section 3.4), which might shed light on evolutionarily beneficial growth strategies (discussed below).

Though IMs derived in this study provided regulatory information for the reconstruction of the TRN of *L. plantarum*, the performance of ICA decomposition was limited by the size of the expression matrix, compared to other ICA-based studies of bacterial transcriptomes (e.g., ICA of *Corynebacterium glutamicum* collected 263 samples from 29 independent projects [38]). Compared to well-studied organisms such as *Escherichia coli*, the amount of existing transcriptomic data of *Lactobacillus plantarum* on NCBI Gene Expression Omnibus (https://www.ncbi.nlm.nih.gov/geo/) [39] is much smaller. Also, due to the lack of operon annotation in *Lactobacillus plantarum*’s genome, motif search for TF binding sites in this study used estimated promoter regions, which lowered the accuracy and might explain why some IMs were uncharacterized. It is also worth noting that the novel regulatory interactions shown by ICA are just “predicted” instead of “confirmed”. To obtain a more valid conclusion, CHiP-seq experiments are needed to confirm those findings in future studies.

With regards to the relationship between secondary and primary metabolism, theoretical models such as Grime’s competitor-stress-ruderal triangle [40][41], Synthetic Chemostat Model[42] and regulatory proteome allocation model [43] all adopted a resource allocation framework to capture the balance between two branches of cellular metabolism. Through the correlations between the activities of identified IMs, this study provided evidence to the theoretical models for secondary metabolism proposed in previous studies by showing the growth strategy of *L. plantarum* that adjusts regulatory activities for different metabolic pathways to react to external stress signals (section 3.4). However, the curated data in this study could not support a significant trade-off relationship between gene expression levels of primary and secondary metabolism. More transcriptomic and proteomic profiling for *L. plantarum* under different growth conditions are needed to quantitatively study the balance between stress and cellular growth.

To conclude, this study provided the *in-silico* TRN reconstruction for *L. plantarum* in a top-down manner and unveiled its growth strategy to balance primary and secondary metabolism with IM activities, in spite of the limitations discussed above. With the growing amount of gene expression data of *L. plantarum* as expected, the quality of IMs derived by ICA will be improved, thus enabling researchers to acquire a better understanding of the underlying rationale of its cellular activities.

## Abbreviation

CPS: capsular polysaccharide
CHiP-seq: chromatin immunoprecipitation followed by sequencing
GLC: glucose
EPS: exopolysaccharide
FOS: fructooligosaccharides
HMO: human milk oligosaccharides
ICA: independent component analysis
IM: iModulon, a set of independently modulated genes
PAC: proanthocyanidin fraction
PCC: Pearson correlation coefficient
TF: transcription factor
TRN: transcriptional regulatory network
XG: xyloglucans.

## Acknowledgements

The authors would like to acknowledge the financial support provided by National Natural Science Foundation of China (project No.32302265; Beijing) and the National Center of Technology Innovation for Dairy (project No. 2023-QNRC-2).

## Author contributions

Sizhe Qiu, Conceptualization, Methodology, Data curation, Data analysis, Writing | Yidi Huang, Data analysis, Writing | Shishun Liang, Data analysis, Writing | Hong Zeng, Writing, Review, Supervision | Aidong Yang, Writing, Review, Supervision

## Data availability statement

The code and data are openly available at https://github.com/SizheQiu/LPiModulons.

